# scMILD: Single-cell Multiple Instance Learning for Sample Classification and Associated Subpopulation Discovery

**DOI:** 10.1101/2025.01.09.632256

**Authors:** Kyeonghun Jeong, Jinwook Choi, Kwangsoo Kim

**Affiliations:** Interdisciplinary Program in Bioengineering, Seoul National University, Seoul, Republic of Korea; Institute of Medical and Biological Engineering, Medical Research Center, Seoul National University, Seoul, Republic of Korea; Department of Transdisciplinary Medicine, Institute of Convergence Medicine with Innovative Technology, Seoul National University Hospital, Seoul, Republic of Korea; Department of Medicine, College of Medicine, Seoul National University, Seoul, Republic of Korea

**Keywords:** Single-cell transcriptomics, Condition-associated cell subpopulations, Multiple Instance Learning, Sample classification

## Abstract

Linking cellular states to clinical phenotypes is a major challenge in single-cell analysis. Here, we present scMILD, a weakly supervised Multiple Instance Learning framework that robustly identifies condition-associated cells using only sample-level labels. After systematically validating scMILD’s accuracy through controlled simulations, we applied it to diverse disease datasets, confirming its ability to retrieve known biological signatures. Building on this, our sample-informed analysis of scMILD-identified monocytes in COVID-19 revealed a temporal transition from an early antiviral to a late stress-response state. Furthermore, in a novel cross-disease application, a model trained on COVID-19 successfully stratified Lupus patients and distinguished shared inflammatory states from disease-specific ones. scMILD thus provides a validated and versatile strategy to dissect cellular heterogeneity, bridging single-cell observations with high-level phenotypes.

## INTRODUCTION

Single-cell transcriptomic data derived from various samples with diverse conditions and phenotypes have been accumulating rapidly^1–3^. However, studies directly analyzing the association between sample conditions/phenotypes and individual cells still need to be completed. The primary analysis of single-cell RNA-seq often focuses on identifying condition/phenotype-specific or associated cell subsets^4–6^. This process typically relies on unsupervised strategies, which can be time-consuming, require extensive knowledge of conditions/phenotypes, and involve subjective judgment from the analyst. These approaches also present challenges for reproducibility and validation, making it difficult to associate individual cells with sample conditions directly.

Recent computational advances have attempted to address these challenges. Deep learning approaches have enabled reference mapping with interpretable gene programs while improving clustering accuracy through various architectural innovations^7^. Several methods have focused on improving cell type identification and clustering accuracy through novel deep learning architectures, particularly addressing the challenges of rare cell populations^8,9^. Other methods have leveraged bulk expression data to identify phenotype-associated subpopulations^10^. While progress has been made in patient classification using single-cell data, current approaches focus on interpretability at the cell type level rather than identifying condition-specific cell states within each type^11^. Moreover, these methods require extensive human intervention or additional data sources, limiting their practical application.

Multiple Instance Learning (MIL) offers a promising framework for addressing these challenges, having demonstrated success in various biological domains, including histopathological image analysis^12–14^. In the context of single-cell transcriptomics, MIL’s bag-based learning paradigm naturally aligns with the hierarchical structure of single-cell data, where samples can be viewed as bags containing multiple cell instances. However, unlike other applications, single-cell transcriptomes present unique challenges due to the need for ground truth labels at the cell level.

To address these limitations, we propose scMILD (Single-cell Multiple Instance Learning for Sample Classification and Associated Subpopulation Discovery), a framework that leverages Multiple Instance Learning to identify condition-associated cell subpopulations while providing quantitative performance metrics through sample-level classification. By treating samples as bags and cells as instances, scMILD offers a systematic approach to bridge sample-level phenotypes with cell-level molecular signatures.

To rigorously validate scMILD, we first demonstrate its technical robustness and the contribution of its key components through systematic simulations and ablation studies. We then benchmark its performance across diverse disease datasets. To showcase scMILD’s power to yield deeper biological insights, we present two distinct advanced applications. First, through a sample-informed analysis of COVID-19 patient data, we reveal dynamic temporal transitions in monocyte states during disease progression. Second, in a novel cross-disease framework, we use a model trained on COVID-19 to dissect cellular heterogeneity in Lupus. Together, these applications demonstrate that scMILD is a versatile tool capable of not only identifying disease-relevant cell populations but also exploring their dynamic changes over time and uncovering shared pathogenic mechanisms across different human diseases.

## RESULTS

### Model Architecture and Training Strategy

The scMILD framework is designed to address the “needle in a haystack” challenge inherent in single-cell analysis, which is governed by two biological assumptions: (1) a sample from a positive condition contains at least one positive-associated cell, and (2) the majority of cells within that positive sample are not associated with the condition. This inherent sparsity means that if these few positive-associated cells can be accurately distinguished from the non-associated background, then assigning meaningful scores and classifying the entire sample becomes a much more direct and robust task. Consequently, scMILD operates on a central modeling premise: that accurate sample classification is best achieved as a consequence of learning a well-structured latent space where positive-associated and non-associated cells are geometrically separable.

This approach addresses a key limitation of standard Multiple Instance Learning (MIL) frameworks such as ABMIL, whose training objective is solely focused on sample-level classification. As a result, a high attention score only signifies importance of an instance for the final prediction, not necessarily its association with the positive condition. For example, a cell with strong “negative” features in a negative sample could receive a high attention score because it helps the model confidently classify the sample as negative. This inherent ambiguity not only complicates the interpretation of attention scores but also means the model has no explicit incentive to organize different cell states within its latent space, failing to guarantee a structured representation suitable for downstream analysis.

To resolve this ambiguity and actively structure the latent space, scMILD introduces a dual-branch architecture centered around a shared Encoder (Figure 1a). This architecture synergistically pursues the macroscopic goal of sample classification and the microscopic goal of refining cellular embeddings. The Sample Branch provides the initial directional signal by using sample-level labels and an attention mechanism to identify potentially positive-associated cells, generating attention scores that serve as pseudo-labels. Guided by these pseudo-labels, the Cell Branch then actively reshapes the latent space. Specifically, it employs a Gaussian Mixture Model (GMM) and an Orthogonal Projection Loss (OPL) to enforce separation, pulling the embeddings of cells with high pseudo-labels closer together while ensuring they are orthogonal to those of cells with low pseudo-labels.

**Figure 1.**
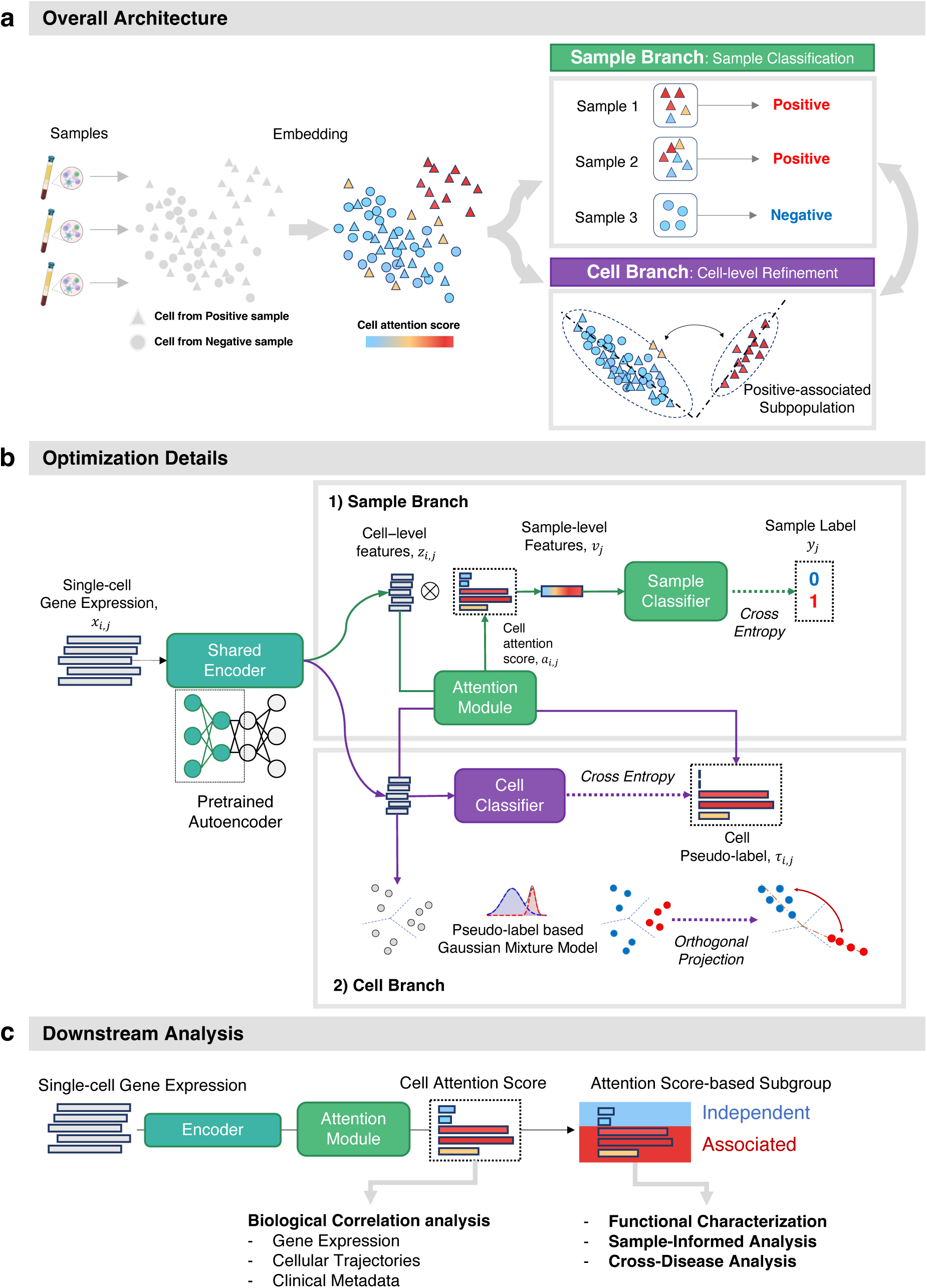
Overview of the scMILD framework. (a) The scMILD model architecture and training process. The Sample Branch performs initial learning using sample-level classification to establish the model foundation, while the Cell Branch refines the model using cell-level prediction with pseudo-labels derived from attention scores. (b) Optimization details of the Sample and Cell Branches. (c) Downstream analysis workflow utilizing cell attention scores for biological correlation analysis, functional characterization, sample-informed analysis and cross-disease analysis.

Through an alternating training process, these two branches cyclically update the shared Encoder (Figure 1b). This synergy ensures that scMILD constructs a well-structured latent space that not only yields high sample classification performance but also provides interpretable and reliable cell-level insights, effectively bridging cellular heterogeneity with high-level sample phenotypes.

### Simulation Study Results

To objectively evaluate scMILD’s performance and practical utility in biological research, we designed controlled simulation studies where ground truth cell labels were available through CRISPR-mediated perturbation (Figure 2a). We first assessed the model’s core capabilities in a single-perturbation setting before testing its robustness in a more complex mixture simulation.

**Figure 2.**
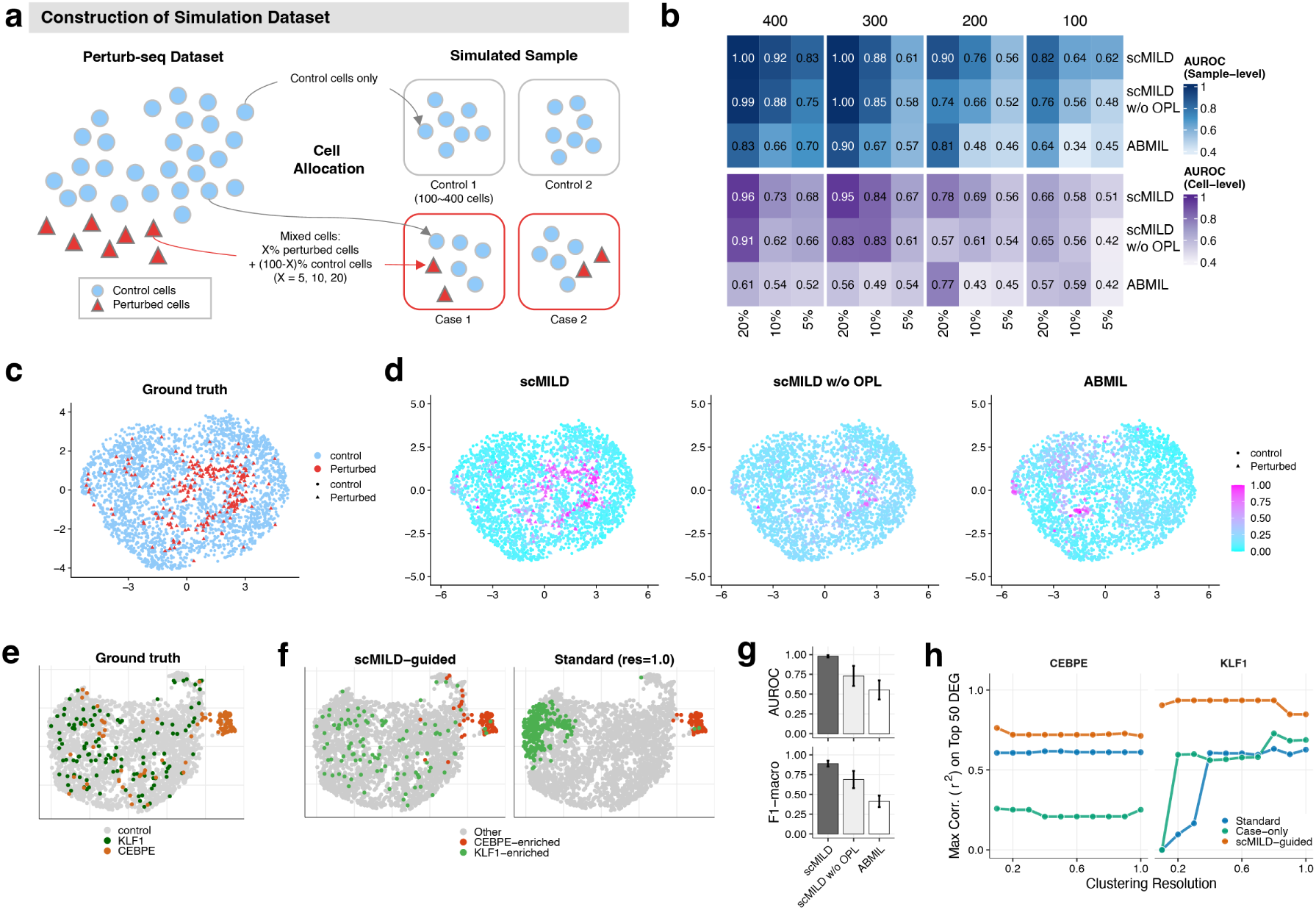
Performance evaluation of scMILD using simulations with single and mixed perturbation types. (a) Schematic of simulated dataset construction from Perturb-seq data, with control (blue circles) and perturbed cells (red triangles) allocated to control and case samples. (b) Heatmap comparing sample and cell classification AUROC for scMILD, scMILD w/o OPL, and ABMIL. Performance is evaluated across varying total cell counts per sample and proportions of perturbed cells in case samples. (c and d) UMAP visualizations of the median-performing test set showing ground truth cell labels (c) and cell attention scores from each model (d), with perturbed cells represented as triangles and control cells as circles. Please define all error bars and statistical tests in the Methods section. (e) UMAP visualization of the mixture simulation dataset, colored by ground truth cell identity (Control, KLF1-perturbed, CEBPE-perturbed). Panels (e-h) show results from this mixture simulation. (f) UMAPs displaying clustering results from scMILD-guided clustering and standard clustering (resolution 1.0). (g) Bar plot comparing sample classification performance (AUROC and F1-macro score) of the three models on the mixture simulation dataset. (h) Coefficient of determination (*r*^2^) from Pearson correlation between the log2 fold-change (log2FC) values of cluster-derived differentially expressed genes (DEGs) and those of the top 50 ground truth DEGs. For each method, the log2FCs from each identified cluster were compared against the ground truth, and the maximum *r*^2^ value achieved is plotted.

#### Performance in Single-Perturbation Simulations

In the single-perturbation setting, analysis of sample classification performance revealed the superior robustness of scMILD across different experimental conditions. To specifically assess the impact of our proposed Orthogonal Projection Loss (OPL), we compared the full scMILD model with an ablated version (scMILD w/o OPL) and a standard attention-based MIL model (ABMIL). With 400 cells and 20% perturbation, scMILD achieved perfect sample classification (AUROC = 1.0), a notable improvement over scMILD w/o OPL (AUROC = 0.989) and ABMIL (AUROC = 0.833) (Figure 2b). Cell-level performance showed similar trends, with scMILD achieving a cell AUROC of 0.956, compared to 0.905 for scMILD w/o OPL and 0.615 for ABMIL. Crucially, the strong correlation between sample and cell-level performance, particularly for the scMILD model, has a significant practical implication (Supplementary Figure S3). Since true cell-level performance cannot be assessed in real-world datasets lacking ground-truth cell labels, a high, empirically measured sample AUROC can serve as a reliable proxy, providing confidence in the biological relevance of the underlying cell attention scores.

We then examined the model’s internal mechanism by analyzing the attention scores assigned to *KLF1*-perturbed and control cells. scMILD demonstrated a more apparent distinction between the two cell groups, achieving higher Kolmogorov-Smirnov statistics and lower overlap coefficients compared to the other models (Supplementary Table S3, Supplementary Figure S1). UMAP visualizations of the median-performing experiment confirmed that scMILD’s cell attention scores (Figure 2d) aligned better with the ground truth cell labels (Figure 2c) than those of the other models. Additional visualizations for experiments with lower cell counts are provided in Supplementary Figure S2.

Finally, we evaluated the practical utility of the identified cell groups for downstream analysis. Using cell attention scores to guide differential expression analysis, scMILD identified substantially more true-positive DEGs with higher recall rates compared to a standard phenotype-based analysis, especially in scenarios with low cell counts where the standard method failed entirely (Supplementary Table S4). This demonstrates scMILD’s ability to effectively capture biological signals even in challenging, low-cell-count datasets.

#### Robust Identification of Heterogeneous Cell Populations in Mixture Simulations

To evaluate scMILD’s performance in more complex scenarios, we extended our framework to a mixture simulation including two distinct perturbed cell types (*KLF1*- and *CEBPE* -perturbed) mixed with control cells (Figure 2e). In this challenging sample classification task, scMILD achieved a mean AUROC of 0.9792 and an F1-macro score of 0.8889, consistently demonstrating superior performance over the ablated version, scMILD w/o OPL (AUROC = 0.7292, F1-macro = 0.6872), and the baseline ABMIL (AUROC = 0.5521, F1-macro = 0.4118) (Figure 2g).

To further evaluate the utility of scMILD for downstream analysis, we compared three clustering strategies: scMILD-guided clustering, standard clustering of all cells, and clustering of only case-derived cells (Case-only). A critical finding was the strong dependency on the resolution parameter exhibited by the standard and Case-only approaches, contrasted with the remarkable stability of scMILD-guided clustering. While the standard methods’ performance varied dramatically with resolution, scMILD maintained consistently high clustering quality across a broad range of resolution values, as measured by Adjusted Rand Index and Adjusted Mutual Information (Supplementary Figure S4d). This stability is a key advantage in real-world applications where optimal parameters cannot be known a priori.

Visualization of the clustering results further illustrated these differences. The scMILD-guided approach effectively separated *KLF1*- and *CEBPE* -perturbed cells into two distinct clusters, which we denote as *KLF1*-enriched and *CEBPE* -enriched (Figure 2f). In contrast, standard and Case-only clustering methods either failed to separate one of the perturbed populations or fragmented them across multiple clusters depending on the chosen resolution (Supplementary Figure S4a-c).

Finally, to assess the biological relevance of the identified clusters, we calculated the coefficient of determination (*r*^2^) from the Pearson correlation between the log2 fold-changes (log2FC) of cluster-based DEG analysis results and the log2FCs of the top 50 ground truth DEGs (Figure 2h). For *KLF1*-perturbed cells, the log2FC values from the scMILD-guided *KLF1*-enriched cluster showed a remarkably high correlation with the ground truth (*r*^2^ = 0.9353), outperforming the best-matching clusters from both Standard (*r*^2^ = 0.6322) and Case-only (*r*^2^ = 0.7288) approaches. For *CEBPE* -perturbed cells, the scMILD-guided approach again achieved the highest correlation (*r*^2^ = 0.7631), substantially outperforming both the Standard (*r*^2^ = 0.6171) and Case-only (*r*^2^ = 0.2587) methods. This robust performance was maintained when correlating against all ground-truth DEGs (Supplementary Figure S4e). Notably, for the Standard and Case-only comparisons, we selected the best-matching cluster from multiple candidates—an ideal scenario impossible in real analysis without prior knowledge. These results confirm that scMILD provides a more robust framework, yielding biologically accurate cell groupings while simultaneously reducing parameter sensitivity and eliminating subjective post-clustering decisions.

### Performance Evaluation on Disease Datasets

To evaluate scMILD’s effectiveness on real-world data, we compared its performance with several other models, including an ablated version of our own model (scMILD w/o OPL) to demonstrate the impact of its key components, as well as established models like ABMIL^12^ and Pro-toCell4P^11^. For a fair comparison between scMILD, scMILD w/o OPL, and ABMIL, we used an identical pre-trained autoencoder architecture, ensuring that performance differences highlight the impact of different architectural components.

We assessed performance using AUROC and F1-macro scores. As shown in Table 1, scMILD demonstrated superior or comparable performance across all datasets and metrics. For instance, in the challenging UC dataset, scMILD substantially outperformed all other models (AU-ROC: 0.9750 vs. 0.9583 for scMILD w/o OPL). This is particularly noteworthy given the dataset’s inflammation-specific cell types, suggesting scMILD’s ability to capture complex disease-specific cellular patterns.

**Table 1.** Performance comparison of scMILD with other models on disease datasets.

| **Metric** | **Dataset** | **scMILD** | **scMILD w/o OPL** | **ABMIL** | **ProtoCell4P** |
| --- | --- | --- | --- | --- | --- |
| **AUROC** | Lupus | **1.0000** | 0.9997 | 0.9968 | **1.0000** |
|  | COVID-19 Infection | **0.9063** | 0.8854 | 0.8646 | 0.8493 |
|  | COVID-19 Hosp. | **0.9562** | 0.9529 | 0.9555 | 0.9549 |
|  | UC | **0.9750** | 0.9583 | 0.8917 | 0.9370 |
| **F1-macro** | Lupus | **0.9801** | 0.9765 | 0.9635 | 0.9742 |
|  | COVID-19 Infection | **0.7769** | 0.7562 | 0.6620 | 0.7097 |
|  | COVID-19 Hosp. | **0.9020** | 0.8880 | 0.8957 | 0.8754 |
|  | UC | **0.9001** | 0.8509 | 0.8514 | 0.5941 |

Notably, the full scMILD model consistently outperformed the scMILD w/o OPL baseline, which in turn outperformed the standard ABMIL model across all datasets. This stepwise improvement demonstrates the effectiveness of scMILD’s architectural design. The performance gain of scMILD w/o OPL over ABMIL highlights the benefit of the dual-branch structure itself. The further improvement of the full scMILD model demonstrates the significant contribution of the Cell Branch’s explicit refinement module, which employs both a Gaussian Mixture Model (GMM) and the Orthogonal Projection Loss (OPL) to structure the cellular embedding space. These results indicate that the components of scMILD work synergistically to achieve robust classification performance. Detailed performance comparisons are provided in Supplementary Table S1.

### Validation of scMILD-identified Condition-associated Cell Subpopulations

To validate the effectiveness of scMILD in identifying condition-associated cell subpopulations, we applied our model to four distinct datasets representing different disease conditions: Lupus, COVID-19 infection, COVID-19 hospitalization, and Ulcerative Colitis. For each dataset, we conducted downstream analysis on the median-performed test set. Our approach involved applying a 2-component Gaussian mixture model based on cell attention scores to divide cells derived from Positive condition samples into two subgroups. The subgroup with higher scores was designated as the condition-associated subgroup, while the other was termed the condition-independent subgroup.

We then focused on identifying and characterizing the condition-associated subgroup, comparing it with known subtypes or markers reported in previous studies. This systematic approach allowed us to evaluate scMILD’s performance across diverse pathological contexts and validate our findings against existing knowledge. Our analysis demonstrates the model’s capability to detect biologically relevant cell subtypes across various disease conditions, as detailed in the following sections.

#### Lupus: Identification of SLE-associated Cell Subtypes

The Lupus dataset comprised 50 samples from healthy individuals and 119 samples from systemic lupus erythematosus (SLE) patients. Figure 3a and b present UMAP visualizations of the dataset, providing insights into cellular heterogeneity and SLE-associated changes. In Figure 3a, we observe distinct clusters corresponding to different cell types, with notable populations of B, T, and monocytes. Figure 3b reveals the distribution of Healthy, SLE-independent, and SLE-associated subgroups within these cell type clusters. Interestingly, the SLE-associated cells are not confined to specific regions but are distributed across multiple cell types, suggesting that SLE-associated changes occur within various cell populations.

**Figure 3.**
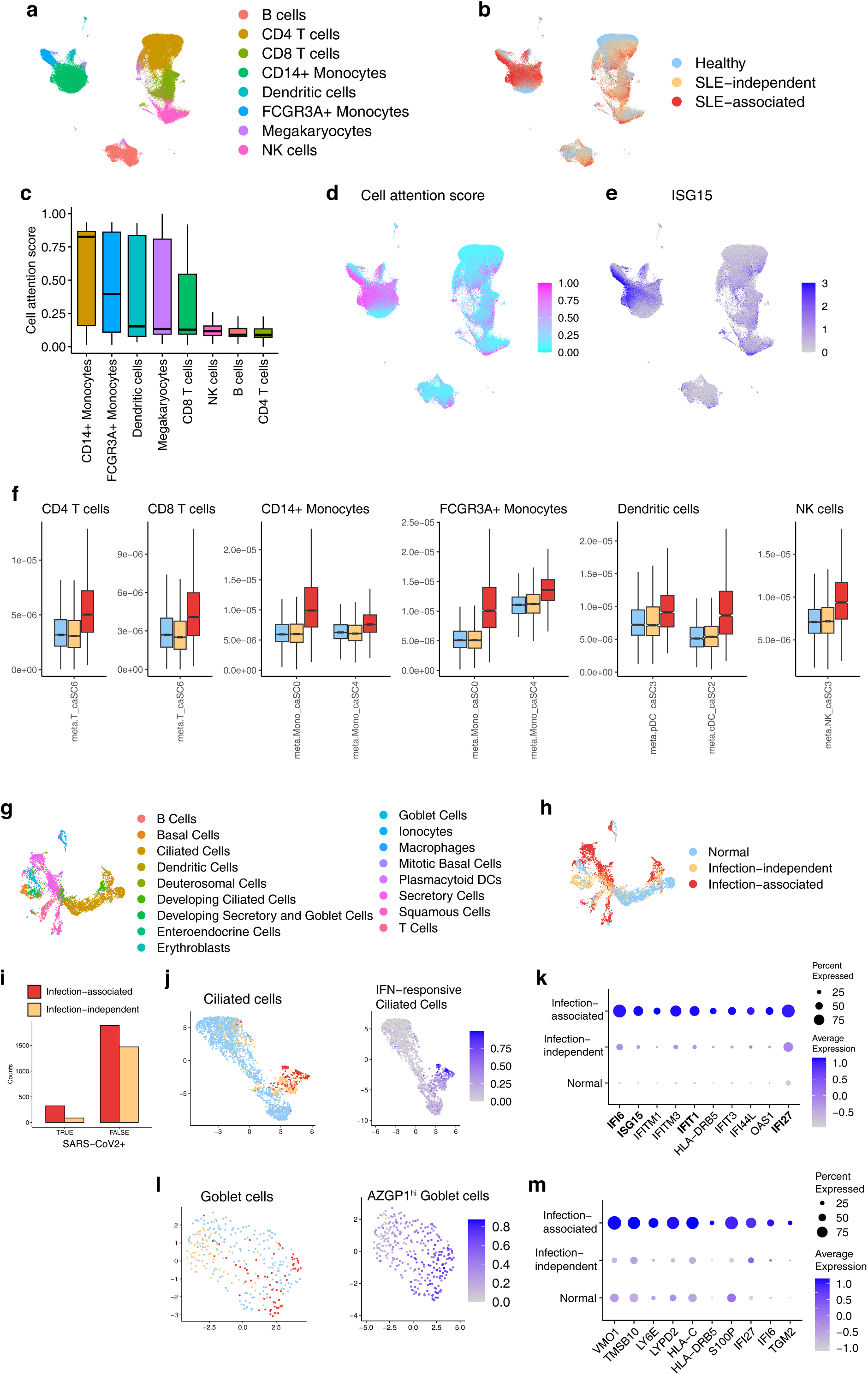
scMILD identifies disease-associated cell subpopulations in Lupus and COVID-19 infection datasets. (a) UMAP visualization of the Lupus dataset, colored by cell type. (b) UMAP visualization of the Lupus dataset, colored by SLE association subgroup (Healthy, SLE-independent, and SLE-associated). (c) Box plot showing the distribution of cell attention scores for each cell type in the Lupus dataset. (d) Feature plot of cell attention scores in the Lupus dataset. (e) Feature plot showing the expression of ISG15 across cells in the Lupus dataset. (f) Box plots comparing meta-feature (ISG-high SLE-expanded subcluster marker gene) expression scores across Healthy, SLE-independent, and SLE-associated subgroups for different cell types in the Lupus dataset. (g) UMAP visualization of the COVID-19 infection dataset, colored by cell type. (h) UMAP visualization of the COVID-19 infection dataset, colored by infection association subgroup (Normal, Infection-independent, and Infection-associated). (i) Bar plot showing the proportion of SARS-CoV2 mRNA-detected cells in each subgroup. (j) UMAP plot of infection association subgroup and Interferon-responsive Ciliated Cell subtype signature scores for Ciliated Cells. (k) Dot plot of top 10 differentially expressed genes in the Infection-associated subgroup of Ciliated Cells. (l) UMAP plot of infection association subgroup and AZGP1-high Goblet cell subtype signature scores. (m) Dot plot of top 10 differentially expressed genes in the Infection-associated subgroup of Goblet Cells. Please define all error bars and statistical tests in the Methods section.

Our analysis revealed that CD14+ Monocytes and FCGR3A+ Monocytes exhibited the highest cell attention scores, consistent with previous research findings on SLE-associated cell types (Figure 3c). This observation aligns with the known involvement of monocytes in SLE pathogenesis.

To further investigate cellular heterogeneity, we employed Gaussian mixture modeling on cell attention scores, independent of cell type, to divide cells from SLE patients into “SLE-associated” and “SLE-independent” subgroups. We then compared our findings with previously reported SLE-associated cell subtypes.

We created a marker gene expression score by leveraging the 100 marker genes for each ISG^hi^ SLE-expanded subcluster reported in a previous study^15^. Notably, the *ISG15* gene, common to all subclusters, showed a high Pearson correlation coefficient of 0.5997 with our cell attention scores (Figure 3d). The feature plot in Figure **??**e further illustrates the expression pattern of *ISG15* across cells in the UMAP space, clearly showing higher expression in regions corresponding to the SLE-associated subgroup.

Furthermore, we compared the expression of meta-features (Top 100 marker genes of ISG^hi^ SLE-expanded subcluster) across Healthy, SLE-independent, and SLE-associated subgroups for different cell types (Figure 3f). The top 100 differentially expressed genes for each ISG^hi^ SLE-expanded subcluster used in this analysis are listed in Supplementary Table S6. The box plots reveal a consistent pattern across all examined cell types: the SLE-associated subgroup demonstrates significantly higher expression of these marker genes than the Healthy and SLE-independent subgroups. This trend is observed across various cell types, including monocytes, dendritic cells, and lymphocytes.

These results validate scMILD’s ability to identify disease-relevant cell subpopulations in SLE, aligning with and extending previous findings in the field. The consistency of our results across different cell types and the clear distinction in meta-feature expression between subgroups underscore the robustness of our approach in capturing disease-associated cellular states.

#### COVID-19 Infection: Detection of Virus-specific Cellular Responses

The COVID-19 infection dataset, which includes measurements of SARS-CoV2 mRNA expression, provided insights into cellular heterogeneity and infection-associated changes. Figure 3g and h present UMAP visualizations of the dataset, revealing distinct cell type clusters and the distribution of Normal, Infection-independent, and Infection-associated subgroups within these clusters.

The Infection-associated cells were distributed across multiple cell types rather than forming a separate cluster. These cells tended to be located towards the periphery of their respective cell type clusters, suggesting that they represent states deviating from typical cellular profiles in normal conditions. Figure 3i quantitatively demonstrates the enrichment of SARS-CoV2 mRNA-detected cells in the infection-associated subgroup across all cell types, supporting the biological relevance of our identified subgroup.

To validate our findings, we compared them with COVID-19 expanded cell subtypes reported by Ziegler et al.^16^. We focused on Ciliated Cells (1571:541, Normal:Infection) and Goblet cells (162:134, Normal:Infection), which had more than 100 cells in both normal and infection samples (Supplementary Table S7).

For Ciliated Cells, we examined the distribution of the Interferon-responsive Ciliated Cell subtype, previously reported as expanded in COVID-19 patients. Using the top 5 DEG markers (*IFI6*, *ISG15*, *IFIT1*, *IFI27*, *MX1*) for this known subtype, we observed a strong correlation between the subtype signature score and our infection-associated subgroup (Figure 3j). The Pearson correlation coefficient between the known subtype signature score and our cell attention score was remarkably high at 0.6348. Moreover, all five known subtype markers were present in the DEGs of our infection-associated subgroup, with four of them (excluding MX1) appearing in the top 10 DEGs (Figure 3k). The dot plot in Figure **??**k clearly shows the upregulation of these interferon-responsive genes in the Infection-associated subgroup compared to the Normal and Infection-independent subgroups.

For Goblet cells, we analyzed the AZGP1^hi^ goblet cell subtype, another COVID-19 expanded cell subtype reported in the original study. Due to the very low p-values of the DEGs provided in the original paper, we selected the top 30 genes to calculate the signature score for this cell subtype. Similar to our findings in Ciliated cells, we observed a strong association between this signature score and our infection-associated subgroup (Figure 3l). The Pearson correlation coefficient between the signature score and our cell attention score was high at 0.6564. Figure 3m shows the top 10 differentially expressed genes in the Infection-associated subgroup of Goblet Cells, highlighting these cells’ distinct gene expression profiles in response to infection. Notably, many of these genes, such as *LY6E*, *IFI6*, and *IFITM3*, are known to be involved in the interferon response, further confirming the biological relevance of our identified subgroups.

These results demonstrate scMILD’s ability to effectively identify virus-specific cellular responses in COVID-19 infection, corroborating previous studies while providing additional granularity in cell subtype identification.

#### COVID-19 Hospitalization: Characterization of Severity-associated Cell Populations

We classified cells into Hosp-associated, Hosp-independent, and Mild subgroups in the COVID-19 Hospitalization PBMC dataset, comprising 254 COVID-19 PBMC samples. Figure 4a and b present UMAP visualizations of the dataset, revealing distinct cell type clusters and precise separation between the hospitalization association subgroups. The UMAP plots demonstrate that certain cell types, mainly monocytes, show a higher proportion of Hosp-associated cells, suggesting their potential role in disease severity.

**Figure 4.**
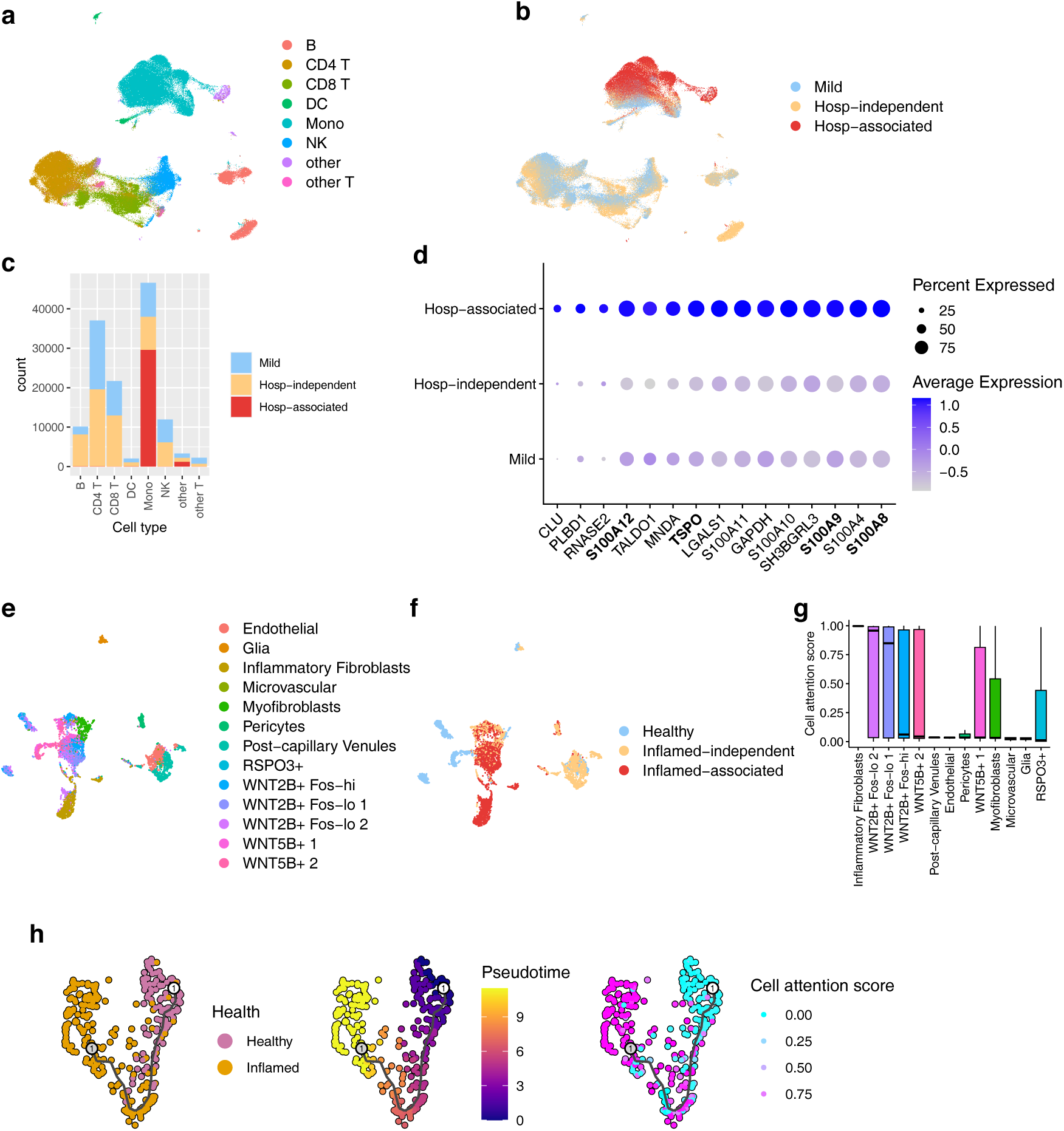
Characterization of scMILD-identified cell states in COVID-19 hospitalization and Ulcerative Colitis. (a) UMAP visualization of the COVID-19 Hospitalization dataset, colored by cell type. (b) UMAP visualization of the dataset, colored by hospitalization association subgroup (Mild, Hosp-independent, and Hosp-associated). (c) Stacked bar plot displaying the number of cells in each subgroup for different cell types. (d) Dot plot showing the expression levels and percentage of cells expressing the top differentially expressed genes in the Hosp-associated CD14+ Monocytes subgroup compared to other subgroups. (e and f) UMAP visualization of the UC test dataset colored by cell type (e) and subgroup (f). (g) Distribution of cell attention scores for each cell type according to sample condition in the UC dataset. (h) Trajectory analysis results for WNT2B+ Fos-lo 2 cells. Each column is colored by sample condition, pseudotime, and cell attention score.

Most cells in the Hosp-associated subgroup were identified as Monocytes at the broad cell type level, with CD14+ monocytes being the predominant cell type at a more specific level. This is clearly illustrated in Figure 4c, where the stacked bar plot shows a higher proportion of Hosp-associated cells in the Mono (Monocyte) category than other cell types.

Our analysis revealed a striking similarity between the Hosp-associated subgroup and the “Dysfunctional CD14 Monocytes” subtype reported in the original study^4^. All marker genes for this subtype (*S100A8*, *S100A12*, *S100A9*, *S100A6*, and *TSPO*) were among the top 15 differentially expressed genes in our Hosp-associated subgroup within the CD14+ monocytes cell type. Figure 4d presents a dot plot of these differentially expressed genes, where the size of the dots represents the percentage of cells expressing the gene and the color intensity indicates the average expression level. The plot clearly shows the upregulation of these marker genes in the Hosp-associated subgroup compared to the Hosp-independent and Mild subgroups, particularly for genes like *S100A8*, *TSPO*, *S100A9*, and *S100A12*.

These results validate scMILD’s ability to identify severity-associated cell populations in COVID-19, aligning closely with previous findings and potentially offering new insights into the cellular basis of disease severity. Identifying these dysfunctional monocytes as a critical feature of severe COVID-19 cases highlights the potential of scMILD in uncovering clinically relevant cellular subpopulations in complex diseases.

#### Ulcerative Colitis: Identification of Inflammation-specific Fibroblasts

In the Ulcerative colitis dataset, we focused on inflammatory fibroblasts, a cell type-specific to the inflamed condition (Figure 4e,f). Our analysis confirmed that scMILD appropriately assigned high cell attention scores to this cell type, validating the model’s ability to recognize disease-specific cell populations (Figure 4g). Interestingly, the WNT2B+ Fos-lo 2 cell type exhibited the second-highest median cell attention score, with a wide distribution. Further investigation through pseudotime analysis revealed a strong correlation between pseudotime and cell attention scores within this cell type, with a Pearson correlation coefficient of 0.6 (Figure 4h). This correlation suggests that the cell attention scores assigned by scMILD may reflect these cells’ developmental or activation trajectory in the context of ulcerative colitis.

These findings corroborate the original study’s identification of inflammatory fibroblasts as a hallmark of inflamed tissue in ulcerative colitis and provide new insights into the potential developmental trajectories of disease-associated cell types.

In summary, across these four diverse disease contexts, scMILD consistently demonstrated its ability to identify condition-associated cell subpopulations that align with and extend previous findings. Our analysis revealed several key strengths of the scMILD approach: 1. Robustness: scMILD effectively identified relevant cell subpopulations across various pathological conditions, from autoimmune diseases to viral infections and inflammatory disorders. 2. Biological relevance: The condition-associated subgroups identified by scMILD showed strong correlations with known disease-specific markers and subtypes reported in previous studies. 3. Novel insights: Besides corroborating existing knowledge, scMILD provided new perspectives on cellular heterogeneity in disease contexts, such as the potential developmental trajectories of disease-associated cell types in ulcerative colitis. 4. Versatility: scMILD’s performance across different tissue types (e.g., PBMCs, nasal epithelium, colon) demonstrates its adaptability to various biological systems. 5. Granularity: The model’s ability to identify condition-associated cells within specific cell types offers a more nuanced understanding of disease processes at the cellular level.

These results demonstrate scMILD’s robustness and potential for discovering novel insights into cellular heterogeneity in various pathological conditions. By leveraging cell attention scores to identify associated subpopulations, scMILD provides a powerful tool for uncovering the cellular basis of complex diseases, potentially leading to new avenues for therapeutic interventions and personalized medicine approaches.

### Sample-Informed Analysis of Cellular States: A COVID-19 Hospitalization Case Study

To investigate the functional heterogeneity within disease-relevant monocytes identified by scMILD, we employed a sample-informed analytical approach focused on high-attention CD14+ monocytes from COVID-19 hospitalized patients. By generating pseudobulk expression profiles from these cells for each sample, we preserved both cellular transcriptional signatures and their sample-level context.

Unsupervised clustering of these pseudobulk profiles revealed five distinct sample groups (Figure 5a), with three clusters displaying characteristic functional signatures.

**Figure 5.**
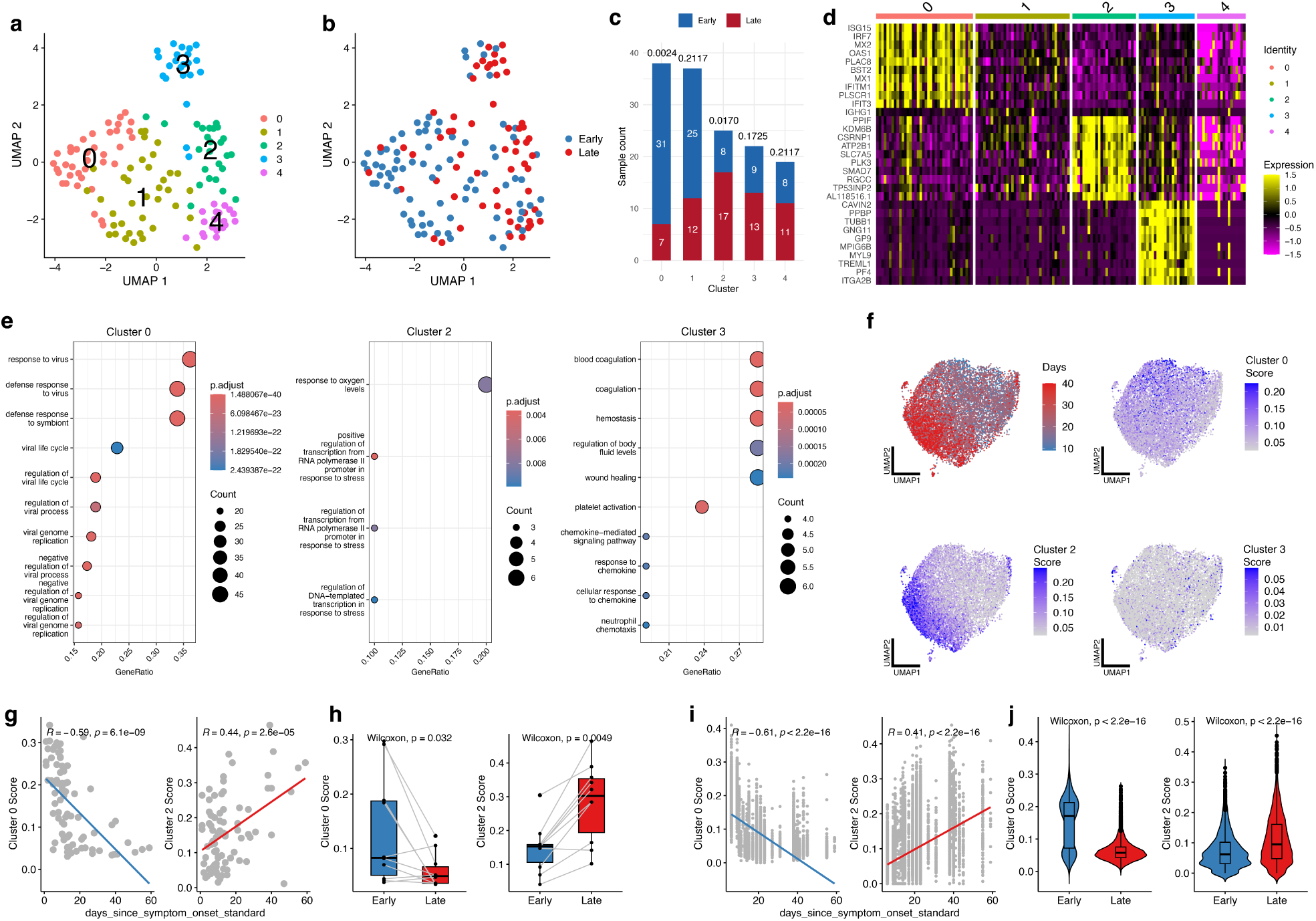
Sample-informed analysis reveals temporal monocyte dynamics in COVID-19 hospitalization. (a) UMAP visualization of pseudobulk profiles from high-attention CD14 monocytes, colored by clustering results showing five distinct sample groups. (b) UMAP visualization of the same profiles colored by time points (Early, Late post-symptom onset). (c) Stacked bar plot showing the distribution of samples collected at early and late across each cluster with Fisher’s exact test p-values indicated. (d) Heatmap displaying the top 10 differentially expressed genes for each pseudobulk cluster. (e) Dot plot of Gene Ontology enrichment analysis results for clusters 0, 2, and 3. (f) UMAP visualization of single cells from the external validation dataset, colored by days post-symptom onset and score distributions for clusters 0, 2, and 3. (g) Scatter plot illustrating the correlation between cluster scores and time since symptom onset at the pseudobulk sample level. (h) Paired box plots comparing cluster 0 and cluster 2 scores between early and late timepoints in longitudinal samples from the same patients. (i) Scatter plot showing the correlation between cluster scores and time since symptom onset at the single-cell level. (j) Violin plots comparing the distribution of cluster 0 and cluster 2 scores between early and late cells. Please define all error bars and statistical tests in the Methods section.

Differential gene expression analysis identified cluster-specific marker genes (Figure 5d). Cluster 0 exhibited elevated expression of interferon-stimulated genes (*ISG15*, *IRF7*, *MX2*, *IFITM1*, *MX1*). Cluster 2 was defined by upregulation of stress response genes (*PPIF*, *KDM6B*, *CSRNP1*, *ATP2B1*, *PLK3*). Cluster 3 showed upregulation of genes associated with platelet activation processes (*CAVIN2*, *PPBP*, *TUBB1*, *GP9*, *PF4*, *ITGA2B*).

Gene Ontology enrichment analysis revealed distinct biological pathways associated with each major cluster (Figure 5e). Cluster 0 showed significant enrichment of antiviral response pathways. Cluster 2 was enriched for response to oxygen levels and stress-induced transcriptional regulation pathways. Cluster 3 demonstrated enrichment of platelet-related processes.

Integration of clinical metadata revealed distinct temporal associations in monocyte transcriptional states (Figure 5b). When samples were colored by time since symptom onset, a separation between early and late samples emerged on the UMAP projection. Quantitative analysis confirmed this temporal association (Figure 5c): the Early Antiviral Response profile (Cluster 0) predominated in samples collected early post-symptom onset (31/38 samples, Fisher’s exact test p=0.0024), while the Stress Response profile (Cluster 2) was significantly associated with samples collected late post-symptom onset (17/25 samples, Fisher’s exact test p=0.0170).

Notably, the temporal dynamics of these transcriptional states partially align with previous observations in independent COVID-19 cohorts. The early interferon response signature we identified in Cluster 0 includes established markers such as *ISG15*, which has been reported by^17^ to show highest expression at early time points with consistent decrease over time. For the late-stage Stress Response signature (Cluster 2), prior research by^18^ reported *PPIF* as one of only two genes showing increased expression at later disease stages, but failed to establish a significant gene set-level association with disease progression.

To validate the reproducibility of these temporal dynamics, we applied our analytical framework to an independent COVID-19 dataset with longitudinal sampling^19^. At the pseudobulk level, our identified molecular signatures showed significant correlations with time since symptom onset (Figure 5g; Cluster 0: r=-0.59; Cluster 2: r=0.44). Paired analysis of longitudinal samples from the same patients further confirmed significant shifts in signature scores over time (Figure 5h; Cluster 0 decreased: p=0.032; Cluster 2 increased: p=0.0049).

Our sample-informed approach demonstrates that these temporal patterns are more robustly captured through gene signature scores than individual markers. The correlation between days since symptom onset and our gene signature scores (Cluster 0: r=-0.61; Cluster 2: r=0.41) substantially exceeded the correlations observed with individual genes like *ISG15* (r=-0.42) and *PPIF* (r=0.18). This suggests that coordinated gene programs, rather than individual markers, better reflect the biological transitions occurring during disease progression.

These results demonstrate a programmatic transition in monocyte functionality during COVID-19 progression from an initial interferon-mediated antiviral state to a subsequent stress adaptation state characterized by response to oxygen levels and stress-induced transcriptional regulation. While previous studies have identified individual components of these responses^17,18^, our sample-informed approach provides a comprehensive view of this cellular state transition at both population and single-cell levels, offering insights into the dynamic nature of immune responses during COVID-19 pathogenesis.

### Cross-Disease Application of scMILD Reveals Shared and Specific Cellular States in COVID-19 and SLE

To explore the cross-disease applicability of scMILD, we performed predictions between COVID-19 and SLE. An scMILD model pre-trained on COVID-19 patient data predicted disease states in an independent SLE cohort with high accuracy (AUROC 0.9012, F1-macro 0.8376), suggesting shared pathological cell states or inflammatory signatures between these distinct diseases.

Conversely, an scMILD model trained on SLE data and applied to COVID-19 patient data showed lower predictive performance (AUROC 0.7176, F1-macro 0.7251), indicating performance asymmetry. This asymmetry may arise because acute COVID-19 inflammatory signatures are potentially more broadly shared than the complex, heterogeneous signatures of chronic

SLE. Furthermore, differences in dataset characteristics, such as the relative homogeneity of acute COVID-19 responses versus the diverse clinical spectrum in SLE, could contribute. These observations underscore the importance of considering training data and disease complexities in cross-disease modeling.

Building on these findings, we investigated single-cell heterogeneity within SLE patients by integratively analyzing cell scores from both the COVID-19-trained (scMILD COVID-19 score) and SLE-trained (scMILD Lupus score) models. Cells were categorized as: ‘Shared’ (high scores from both models), ‘Lupus-Dominant’ (LD; high Lupus score, low COVID-19 score), ‘COVID19-Dominant’ (CD; high COVID-19 score, low Lupus score), or ‘Control’ (low scores from both models or healthy control cells) (Figure 6a).

**Figure 6.**
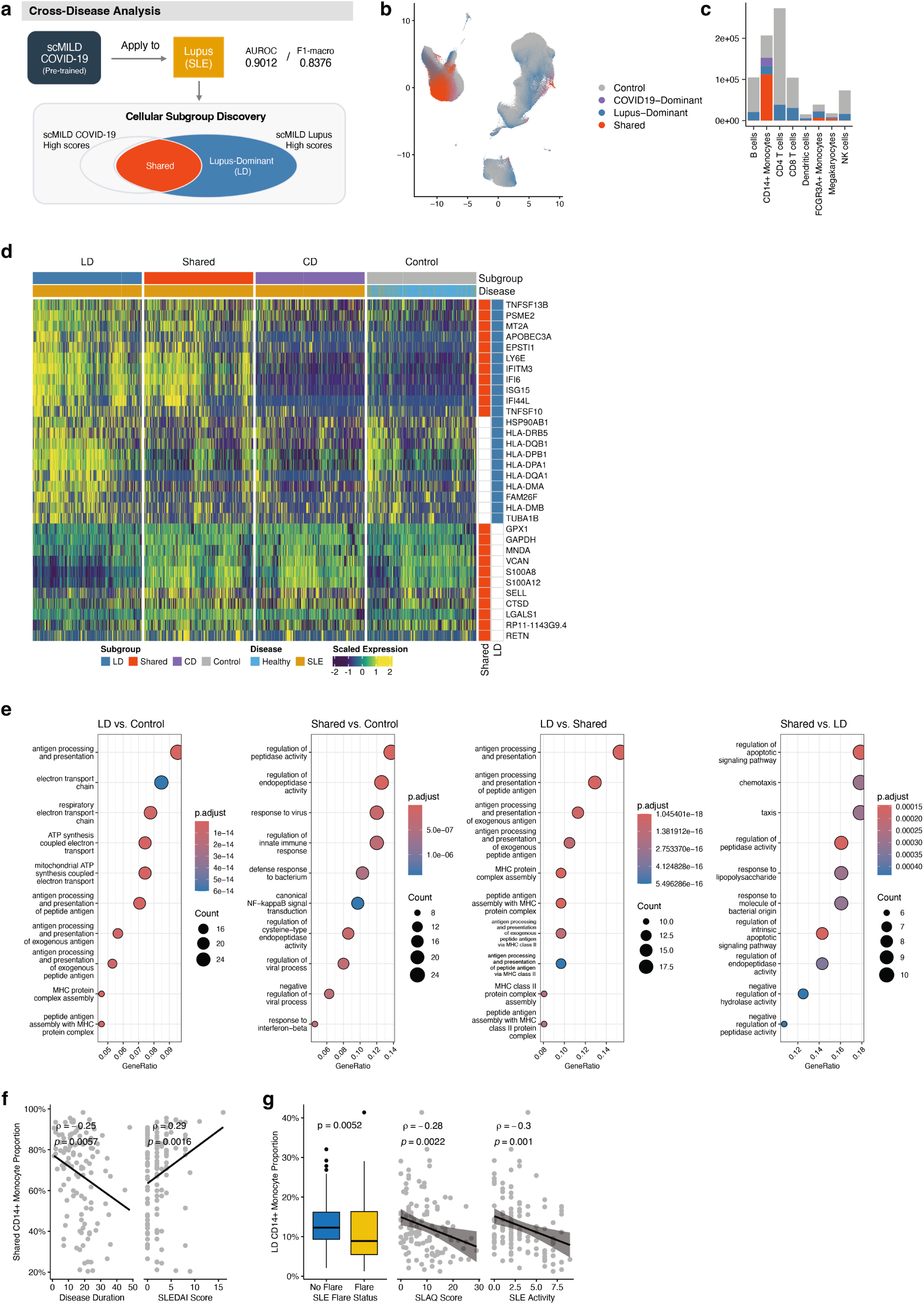
Cross-Disease Application of scMILD Reveals Shared and Specific Cellular States in COVID-19 and SLE. (b) Schematic of the cross-disease analysis workflow. An scMILD model pre-trained on COVID-19 data is applied to the SLE dataset. Cells are then categorized into Shared, Lupus-Dominant (LD), COVID19-Dominant, and Control subgroups based on their attention scores. (c) UMAP visualization of all cells from the SLE dataset, colored by the cross-disease subgroups defined in (a). (d) Stacked bar plot showing the cellular composition of each cross-disease subgroup across major cell types. (e) Heatmap showing scaled expression of the union of top 10 DEGs from the Shared vs. Control and LD vs. Control comparisons. The heatmap includes 1,000 subsampled cells per group, with rows annotated to highlight genes specific to Shared or LD signatures. (f) Dot plot displaying the top 10 enriched Gene Ontology (GO) Biological Process terms for differentially expressed genes (DEGs) from comparisons between Shared, Lupus-Dominant (LD), and Control CD14+ Monocytes. (g) Scatter plots showing the correlation between the proportion of Shared CD14+ Monocytes and clinical variables, including disease duration and SLEDAI score. (h) Clinical associations for the proportion of Lupus-Dominant (LD) CD14+ Monocytes. A box plot compares proportions by SLE flare status, and scatter plots show correlations with SLAQ score and overall SLE activity.

Analysis of these cellular subgroup distributions across all cell types (Figure 6b, 6c) revealed ‘Shared’ cells were most prominent within CD14+ monocytes (54.1%). This aligns with previous findings of high scMILD scores in CD14+ monocytes from severe COVID-19 patients suggesting a key role for these cells in common pathogenic mechanisms. Subsequent analyses focused on the molecular features of ‘Shared’ and ‘LD’ CD14+ monocyte subpopulations in SLE.

Differential gene expression (DEG) analysis comparing ‘Shared’, ‘LD’, and ‘Control’ CD14+ monocytes (Figure 6d) showed that interferon-response genes (*ISG15*, *IFI6*, *IFITM3*) were commonly upregulated in both ‘Shared’ and ‘LD’ groups versus ‘Control’ cells, indicating activated interferon-mediated inflammation. Direct comparison revealed distinct signatures: ‘LD’ CD14+ monocytes had significantly higher expression of MHC class II-related genes (*HLA-DQA1, HLA-DPB1*), while ‘Shared’ CD14+ monocytes were characterized by high expression of previously reported dysfunctional monocyte markers from COVID-19 (*S100A8/A9/A12, TSPO*).

GO enrichment analysis (Figure 6e) showed ‘LD’ CD14+ monocytes were primarily enriched for “antigen processing and presentation” GO biological process (BP) terms compared to both ‘Control’ and ‘Shared’ groups, suggesting enhanced antigen presentation capacity. ‘Shared’ CD14+ monocytes, versus ‘Control’, were enriched for “innate immune response,” “response to virus,” and “response to interferon-beta.” Compared to ‘LD’ cells, ‘Shared’ cells showed enrichment for “regulation of apoptotic signaling pathway” and “regulation of peptidase activity,” implying involvement in general inflammatory and specific stress responses. These results molecularly define heterogeneous CD14+ monocyte subpopulations in SLE, exhibiting both COVID-19-shared inflammatory states and lupus-specific immunological features.

To evaluate the clinical significance of these ‘Shared’ and ‘LD’ CD14+ monocyte subtypes in SLE, we correlated their patient-specific proportions with lupus clinical parameters using Spearman correlation and Wilcoxon rank-sum tests (Figures 6f, Supplementary Table S12). The ‘Shared’ CD14+ monocyte proportion positively correlated with SLE Disease Activity Index (SLEDAI) scores (*ρ* = 0.29, FDR < 0.05) and negatively with disease duration (*ρ* = −0.25, FDR < 0.05). This suggests the ‘Shared’ subtype is associated with current disease activity and may be more prominent in earlier or active disease phases.

Conversely, the ‘LD’ CD14+ monocyte proportion was significantly lower in patients with active SLE flares versus non-flare states (Wilcoxon p = 0.0052, Figure 6g). It also negatively correlated with the Systemic Lupus Activity Questionnaire (SLAQ) score (*ρ* = -0.28, FDR <0.05) and overall SLE activity (*ρ* = -0.30, FDR <0.05). These findings suggest that a relative increase in ‘LD’ CD14+ monocytes may associate with disease stability or lower patient-perceived symptoms, potentially linking to their enhanced antigen presentation capacity and reflecting a chronic or immunomodulatory state.

Collectively, these clinical associations indicate that ‘Shared’ and ‘LD’ CD14+ monocyte subtypes are linked to distinct clinical features and disease activity patterns in SLE. This highlights their potential beyond mere cytological distinctions, offering value for understanding patient heterogeneity, assessing disease status, and exploring biomarkers for personalized treatment strategies.

## DISCUSSION

scMILD demonstrated robust performance across diverse single-cell RNA-seq datasets, highlighting its potential for broad application in various biological contexts. The model’s effectiveness stems from its dual-branch architecture, where the Sample Branch performs robust sample-level classification while the Cell Branch identifies condition-associated cell subpopulations through enhanced representation learning. Unlike conventional unsupervised clustering approaches that require extensive knowledge and subjective judgment, scMILD systematically leverages sample-level labels to identify condition-associated cellular populations, consistently outperforming state-of-the-art models such as ABMIL, and ProtoCell4P.

Our mixture simulation studies revealed a significant advantage of scMILD in addressing parameter sensitivity issues that plague standard clustering methods. While conventional approaches require extensive parameter tuning to optimally identify different cell types, scMILD maintained consistent high performance across a broad range of resolution values. This stability is particularly valuable in real-world applications where optimal parameters cannot be determined a priori. Furthermore, scMILD demonstrated superior ability to detect differentially expressed genes compared to phenotype-based analysis, particularly in datasets with limited cell numbers where conventional approaches failed completely.

The utility of scMILD in dissecting cellular heterogeneity is further exemplified by its application in complex disease scenarios. For instance, the sample-informed analysis in COVID-19 hospitalization revealed temporal monocyte transitions, with gene signatures robustly tracking disease progression. Extending this, our cross-disease analysis between COVID-19 and SLE demonstrated scMILD’s capacity to identify both shared inflammatory states and disease-specific cellular characteristics within SLE patients. Crucially, scMILD linked these distinct monocytic states to different clinical trajectories and disease activity levels in SLE, showcasing its power to define clinically relevant cellular heterogeneity by integrating single-cell data with sample-level phenotypes across different pathological contexts. This ability to parse complex cellular landscapes and connect them to clinical outcomes offers a significant advantage over methods that identify cellular changes without direct phenotypic linkage.

This study presents scMILD as a significant contribution to single-cell transcriptomics by establishing a systematic, reproducible method for connecting sample conditions to specific cell subpopulations. CRISPR-perturbation simulation studies definitively demonstrated scMILD’s ability to accurately identify condition-associated cells in both single and mixed perturbation scenarios. The model’s excellent performance, even with limited cell numbers and reduced parameter sensitivity, enhances its practicality. Notably, across diverse disease datasets including Lupus, COVID-19, and Ulcerative Colitis, scMILD not only consistently outperformed state-of-the-art models but also identified condition-associated cell subpopulations that aligned with findings from original studies. These capabilities offer new possibilities for understanding disease-relevant cellular states and monitoring disease progression, ultimately advancing our understanding of cellular heterogeneity and its role in disease mechanisms.

### Limitations of the study

Despite these strengths, several limitations should be acknowledged. First, the current implementation focuses on binary classification problems, requiring extension for more complex multi-class scenarios. Second, the model’s performance depends on the accuracy of sample labels, where incorrect labeling could lead to inaccurate cell-level predictions. Third, interpretability of the features driving cell-level attention scores remains limited, potentially constraining deeper insights into biological mechanisms. Finally, the model identifies condition-associated cells regardless of their annotated type, a strength for discovering novel or cross-type cellular states. A limitation, however, is that the model’s global optimization strategy does not guarantee the identification of all relevant changes within every distinct cell type, especially for subtle alterations. Therefore, a complete assessment of cell-type-specific responses requires dedicated post-hoc integration of the model’s outputs with cell type labels.

## RESOURCE AVAILABILITY

### Lead contact

Requests for further information and resources should be directed to and will be fulfilled by the lead contact, Kwangsoo Kim.

### Materials availability

This study did not generate new materials.

### Data and code availability

- The publicly available datasets used in this study can be accessed from their original repositories. The Lupus dataset is available on GitHub (https://github.com/yelabucsf/ lupus_1M_cells_clean). The COVID-19 Infection dataset is available through the Single Cell Portal under accession code SCP1289. The COVID-19 Hospitalization dataset (Su et al.) can be accessed via the Fred Hutch COVID-19 Atlas (https://atlas.fredhutch. org/fredhutch/covid/). The Ulcerative Colitis dataset is available on GitHub (https://github.com/cssmillie/ulcerative_colitis). The Perturb-seq data used for generating the simulation dataset is available on Zenodo (DOI: 10.5281/zenodo.7041849). Any other data reported in this paper will be shared by the lead contact upon reasonable request.
- All original code developed for this study, including for data analysis and model implementation, has been deposited on GitHub and is publicly available at https://github.com/Khreat0205/scMILD.

## ACKNOWLEDGMENTS

The authors thank the anonymous reviewers for their valuable suggestions. This work was funded by Korea National Institute of Health (KNIH) via grant (No. 2024-ER-0801-01). The authors thank all members of the lab for their support.

## AUTHOR CONTRIBUTIONS

Conceptualization, K.K., J.C.; Methodology, K.J., K.K.; Software, K.J.; Validation, K.J.; Investigation, K.J.; Data Curation, K.J.; Writing – Original Draft, J.C.; Writing – Review & Editing, K.K., J.C.; Supervision, K.K., J.C.

## DECLARATION OF INTERESTS

The authors declare no competing interests.

## DECLARATION OF GENERATIVE AI AND AI-ASSISTED TECHNOLOGIES

During the preparation of this work the authors used Claude (Anthropic) in order to improve language and readability. After using this tool, the authors reviewed and edited the content as needed and take full responsibility for the content of the publication.

## SUPPLEMENTAL INFORMATION INDEX

Figure S1-S4 and their legends in a PDF

Table S1. Performance comparison between scMILD and the baseline model across eight experimental runs.

Table S2. Hyperparameters used for the scMILD model across all datasets.

Table S3. Comparison of attention score separability metrics across simulation settings.

Table S4. Comparison of DEG identification performance on single-perturbation simulation dataset.

Table S5. Comparison of biological relevance of identified clusters in the mixture simulation.

Table S6. Top 100 differentially expressed genes (DEGs) for ISG^hi^ SLE-expanded subclusters.

Table S7. Cell type counts in the COVID-19 infection test dataset by sample phenotype.

Table S8. Distribution of SARS-CoV2 positive cells across subgroups in the COVID-19 infection dataset.

Table S9. Differentially expressed genes in Ciliated cells from the COVID-19 infection test dataset.

Table S10. Differentially expressed genes in Goblet cells from the COVID-19 infection test dataset.

Table S11. Differentially expressed genes in CD14+ Monocytes from the COVID-19 Hospitalization test dataset.

Table S12. Correlation between CD14+ monocyte subtype proportions and clinical variables in SLE.

## STAR METHODS

### Key resources table

(Please see the separate Key Resources Table file for details of all key reagents, software, and resources.)

### Method details

#### Data Acquisition and Preprocessing

##### Simulation Datasets

To systematically evaluate model performance under controlled settings, we designed simulation studies using preprocessed single-cell RNA sequencing data from a CRISPRa-based gene over-expression experiment^20^. The data, containing control cells and cells with specific transcription factor perturbations, was preprocessed and filtered to include only cells with good coverage as described previously^21^. For our primary simulation, we utilized *KLF1*-perturbed cells (1,843 cells) as ground truth condition-specific cells. To simulate more complex scenarios, a mixture simulation dataset was constructed by incorporating both *KLF1*- and *CEBPE* -perturbed cells (1,172 cells). For all simulation datasets, we identified 2,000 highly variable genes (HVGs) for feature selection using scanpy^22^. The detailed design of the simulation experiments is described in the ”Simulation Study Design” subsection below.

##### Disease Datasets

We evaluated scMILD on four public single-cell RNA-seq datasets representing various disease conditions. These include a Lupus dataset (169 samples, 834,096 cells)^23^, a COVID-19 nasal swab dataset (Infection; 50 samples, 26,947 cells)^16^, a COVID-19 peripheral blood mononuclear cell (PBMC) dataset (Hospitalization; 254 samples, 515,141 cells)^4^, and an Ulcerative Colitis (UC) dataset (42 samples, 18,725 cells)^6^. All COVID-19 datasets used in this study (Infection, Hospitalization) were obtained from pre-processed data curated in Tian et al., 2022^24^. For feature selection, we selected 3,000 HVGs for the Lupus, COVID-19 Infection, and COVID-19 Hospitalization datasets, and 2,000 HVGs for the UC dataset. For the COVID-19 Infection dataset, an additional filter was applied to retain genes expressed in at least five cells. Raw counts were used as model input for all datasets.

##### External Validation Dataset for Temporal Analysis

For validation of our sample-informed temporal analysis, we employed an independent longitudinal PBMC dataset from COVID-19 patients^19^. This dataset contains 95 samples from 81 patients (12 non-hospitalized, 83 hospitalized) and includes 27 longitudinal samples from 13 patients, enabling the validation of temporal dynamics. Data preprocessing for this validation set followed the same HVG selection protocol as our primary COVID-19 Hospitalization dataset.

#### The scMILD Framework

scMILD is a weakly supervised deep learning framework based on Multiple Instance Learning (MIL) designed to classify samples and identify condition-associated cell subpopulations from single-cell transcriptomic data. It comprises a shared Encoder and two specialized branches: a Sample Branch and a Cell Branch.

##### Encoder for Cellular Representation

The encoder, an autoencoder-based neural network, learns a low-dimensional latent representation of each cell’s gene expression. It takes a raw gene count vector *x_i,j_* for cell *i* in sample *j* and maps it to a latent vector *z_i,j_* = *f*_enc_(*x_i,j_*), where *z_i,j_*∈ ℛ*^k^*. The decoder part of the autoencoder is used to reconstruct the gene expression profile. The model is trained to minimize the negative log-likelihood of a negative binomial (NB) distribution, which is well-suited for sparse, over-dispersed single-cell RNA-seq data. The NB loss is defined as:

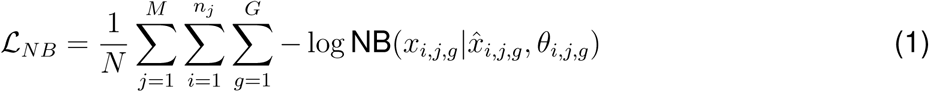

where *N* is the total number of cells, *M* is the number of samples, *n_j_*is the number of cells in sample *j*, and *x̂* and *θ* are the mean and dispersion parameters of the NB distribution output by the decoder. After pretraining, the trained encoder *f*_enc_ is used as a shared feature extractor for both the Sample and Cell branches.

##### Attention Module for Cell Weighting

An attention module, adapted from Ilse et al., 2018^12^, is used to compute an attention score *a_i,j_* ∈ [0, 1] for each cell, reflecting its importance for the sample’s classification. The score is calculated from the latent representation *z_i,j_* as:

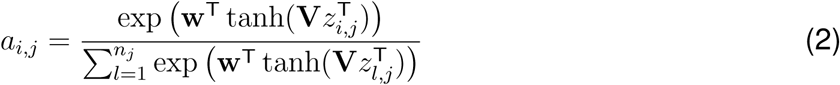

where **V** and **w** are learnable parameters. The attention scores are normalized to sum to 1 within each sample.

##### Sample Branch for Sample-Level Classification

The Sample Branch aggregates cellular information to predict a sample-level label. A sample-level feature vector *v_j_* is computed by taking an attention-weighted sum of the cell latent representations: 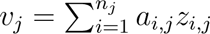. This vector is then fed into a sample classifier 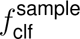 to predict the sample probability 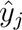. The branch is trained by minimizing the standard binary cross-entropy loss 𝓛_sample_ between the predicted probabilities *ŷ_j_*and the true sample labels *y_j_*:

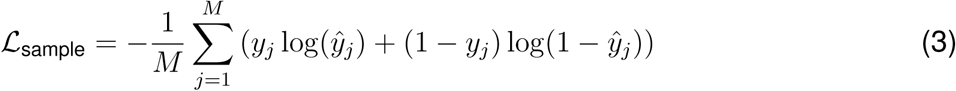

##### Cell Branch for Cell-State Refinement

The Cell Branch aims to refine the cellular representations by performing cell-level predictions. It uses the attention scores from the Sample Branch to generate pseudo-labels for cells: *τ_i,j_* = *a_i,j_* if the sample *j* is positive (*y_j_* = 1), and *τ_i,j_* = 0 if it is negative (*y_j_* = 0). A cell classifier 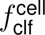 then predicts a cell-level probability 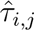 from the latent vector *z_i,j_*. The training objective for this branch combines two loss functions. First, a weighted binary cross-entropy loss, 𝓛_WCE_, is used to handle the imbalance in pseudo-labels:

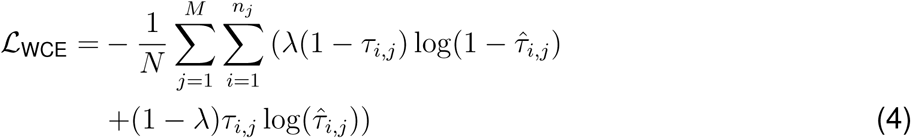

where *λ* is a weight to balance the contribution of positive and negative pseudo-labels. Second, an orthogonal projection loss (OPL), 𝓛_OPL_, is incorporated to enhance the separability of cellular states within positive samples^25^. The OPL encourages embeddings of cells with high pseudo-labels (*C*_high_) to be similar to each other, while being orthogonal to embeddings of cells with low pseudo-labels (*C*_low_). This is achieved by first clustering cells in positive samples into high- and low-attention groups using a Gaussian Mixture Model, and then optimizing the cosine similarity within and between these groups:

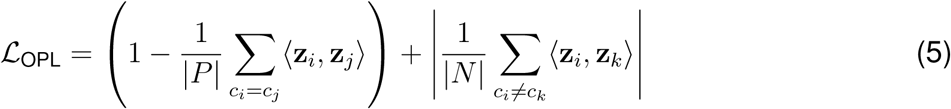

where *c_i_* represents the cluster label (high or low) for cell *i*, |*P* | and |*N* | are the numbers of same-cluster and different-cluster pairs, respectively, and ⟨·, ·⟩ is the cosine similarity operator. The total loss for the Cell Branch is 𝓛_cell_ = 𝓛_WCE_ + *γ*𝓛_OPL_, where *γ* is a hyperparameter.

#### Model Training Procedure

The training of scMILD is performed in two stages.

##### Stage 1: Pretraining the Autoencoder

The autoencoder is first pretrained on all cells from the dataset to learn robust and generalizable cellular representations by minimizing the negative binomial reconstruction loss (𝓛*_NB_*). Key hyperparameters for this stage include the encoder layer dimensions (512, 256, 128), a learning rate of 0.001, and a maximum of 250 epochs with an early stopping patience of 15. The batch size was set to 128.

##### Stage 2: Alternating Optimization of Sample and Cell Branches

After pretraining, the main scMILD model is trained by alternating between optimizing the Sample Branch and the Cell Branch.

1. **Sample Branch Optimization**: The parameters of the encoder, attention module, and sample classifier are updated to minimize the sample-level classification loss (𝓛_sample_).
2. **Cell Branch Optimization**: The parameters of the encoder and cell classifier are updated to minimize the combined cell-level loss (𝓛_cell_), using the attention scores from the latest Sample Branch as pseudo-labels.

The ratio of optimization steps between the two branches is a hyperparameter that varies by dataset (e.g., 1 for most, 5 for the COVID-19 Hospitalization dataset). Optimization is performed using the Adam optimizer with specific learning rates, dimensions, and early stopping criteria detailed in Supplementary Table S2.

#### Simulation Study Design

To assess scMILD’s performance under controlled settings, we constructed binary classification datasets from the simulation data. Each dataset comprised two groups of 14 samples each: control samples (containing only control cells) and case samples (containing a mixture of control and perturbed cells). The number of cells per sample was varied (400, 300, 200, 100), and the proportion of perturbed cells in case samples was also varied (20%, 10%, 5%). For the mixture simulation, case samples contained 10% *KLF1*-perturbed and 10% *CEBPE* -perturbed cells. This design enabled a rigorous evaluation of both sample classification accuracy and cell-level identification capability under diverse conditions.

#### Downstream Analysis of scMILD Outputs

##### Cell Stratification and Condition-Association Grouping

To systematically analyze cell populations based on their relevance to the sample condition, we stratified cells from the test set using the attention scores generated by scMILD’s attention module. For each dataset, we applied a 2-component Gaussian Mixture Model (GMM) to the distribution of attention scores. This partitioned cells into high-attention and low-attention clusters. Based on this, cells from positive-condition samples (e.g., Infected, Inflamed, or Hospitalized) were categorized into two analytical groups:

- **Condition-associated group**: Cells from positive samples belonging to the high-attention GMM cluster.
- **Condition-independent group**: Cells from positive samples belonging to the low-attention GMM cluster.

Cells from negative-condition samples were analyzed separately as a baseline control group. This stratification strategy enabled a focused comparison between cellular states within positive samples.

##### Calculation of Known Subcluster/Subtype Signature Scores

To validate the biological relevance of our identified cell subpopulations, we calculated gene signature scores for known cell subtypes or subclusters reported in the original studies. For the Lupus dataset, we computed meta-feature scores for ISG^hi^ SLE-expanded subclusters using the top 100 marker genes from Nehar-Belaid et al., 2020^15^ with the MetaFeature function in Seurat^26^. For the COVID-19 infection dataset, we used the UCell package^27^ to calculate scores for known subtypes based on their top 5 marker genes as identified in Ziegler et al., 2021^16^.

##### Pseudotime Analysis

To investigate cellular trajectories in the Ulcerative Colitis dataset, we performed pseudotime analysis using Monocle3^28^ on the WNT2B+ Fos-lo 2 cell population. The trajectory graph was learned using the learn graph function with default parameters. The root of the trajectory was set to the principal node with the highest proportion of healthy cells or the lowest mean cell attention score.

##### Sample-Informed Analysis of Cellular States

To investigate functional heterogeneity within condition-associated cells while preserving sample context, we developed a sample-informed analytical workflow. This approach was applied to the COVID-19 Hospitalization dataset, focusing on high-attention CD14+ monocytes. We first generated a pseudobulk expression profile for these cells for each sample using Seurat’s AggregateExpression function. Unsupervised clustering was then performed on these pseudobulk profiles to identify distinct sample groups. Functional interpretation of these groups was carried out through differential expression and gene ontology enrichment analysis.

### Quantification and statistical analysis

#### Experimental Design and Model Evaluation

All experiments were designed to ensure robust and reproducible evaluation of model performance. Each dataset was systematically partitioned into training (50%), validation (25%), and test (25%) sets. To ensure statistical robustness, we conducted eight independent runs for each experimental setting and implemented early stopping based on validation metrics.

Our comparative analysis employed a hierarchical approach to systematically dissect the contributions of scMILD’s components. We used a standard attention-based MIL framework (ABMIL^12^) as the foundational baseline. Building upon this, ‘scMILD w/o OPL; incorporates our dual-branch architecture but lacks the final Orthogonal Projection Loss, allowing us to isolate the impact of the dual-branch structure. The full scMILD model then adds the OPL. This tiered comparison enables a clear, stepwise evaluation of each of our architectural innovations. To ensure a fair comparison, these models were provided with identical preprocessed data and utilized the same pretrained autoencoder architecture. We also compared scMILD with other relevant models where applicable.

#### Performance Metrics

Sample-level classification performance was primarily assessed using the area under the receiver operating characteristic curve (AUROC) and the macro-averaged F1-score. For the simulation studies, where ground truth cell labels were available, cell-level identification performance was also measured by the AUROC of cell attention scores against the true cell labels. Clustering performance in the mixture simulation was quantified using the adjusted Rand index (ARI) and adjusted mutual information (AMI). For evaluating the separation of attention score distributions between perturbed and control cells in simulations, Kolmogorov-Smirnov (KS) statistics and overlap coefficients were calculated.

#### Statistical Analysis

All statistical analyses were performed using R (v4.3.1) or Python (v3.8.18).

##### Differential Gene Expression (DEG) Analysis

DEGs between specified cell groups were identified using the FindMarkers or FindAllMarkers functions in the Seurat R package^26^, which implements a Wilcoxon rank-sum test by default. A gene was considered differentially expressed if the Bonferroni-corrected adjusted p-value was less than 0.01 and the absolute log2 fold-change was greater than 0.25, unless otherwise specified. For the sample-informed analysis, DEG analysis on pseudobulk profiles was performed using the DESeq2 package^29^ with an adjusted p-value cutoff of 0.01.

##### Gene Ontology (GO) Enrichment Analysis

Functional enrichment analysis of DEG lists was performed using the clusterProfiler R package^30^. GO terms in the ”Biological Process” ontology were considered significantly enriched if the associated p-value and q-value were less than 0.05.

##### Correlation and Association Tests

Correlations between variables, such as signature scores and pseudotime, were assessed using Pearson or Spearman correlation coefficients, as specified in the text. Associations between categorical variables, such as sample cluster membership and time points, were evaluated using Fisher’s exact test. For paired comparisons of longitudinal samples, the Wilcoxon signed-rank test was used. For comparisons between two independent groups, the Wilcoxon rank-sum test was used. Statistical significance for all tests was generally defined as p <0.05, unless otherwise noted.

## Additional resources

## Notes

https://github.com/Khreat0205/scMILD

### Competing Interest Statement

The authors have declared no competing interest.

### Summary of Updates

Major additions: New "Sample-Informed Analysis" section and "Cross-Disease Application" section added ; Figures 5 and 6 added to demonstrate sample-informed analysis and cross-disease validation results. Structural changes: Manuscript reformatted to comply with journal submission requirements. Technical improvements: Supplementary materials updated with additional methodological details and performance metrics. Note: This revision represents the version submitted for peer review. Core methodology and main findings remain consistent with the original preprint.

