## Supplementary Figures for "scMILD: Single-cell Multiple Instance Learning for Sample Classification and Associated Subpopulation Discovery"

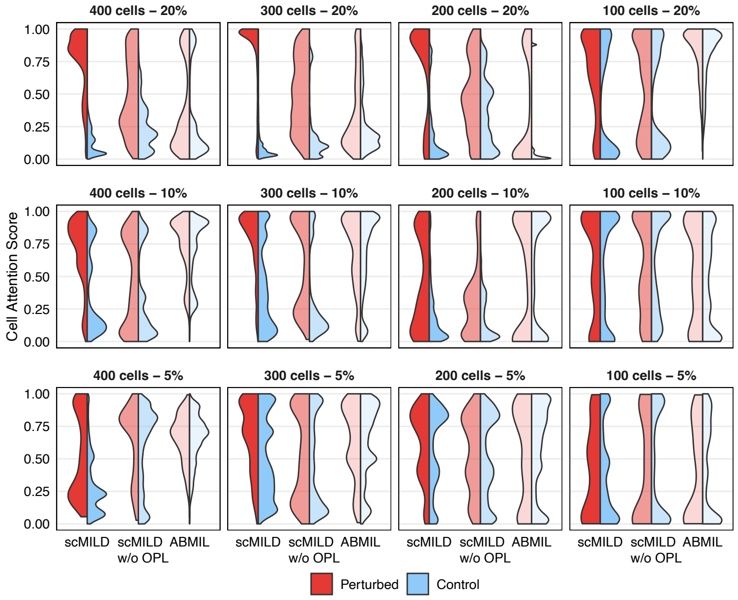


**Supplementary Figure S1.** Split violin plots of cell attention score distributions from scMILD, scMILD w/o OPL, and ABMIL models for perturbed (red) and control (blue) cells across varying total cell numbers (400, 300, 200, and 100 cells per sample).

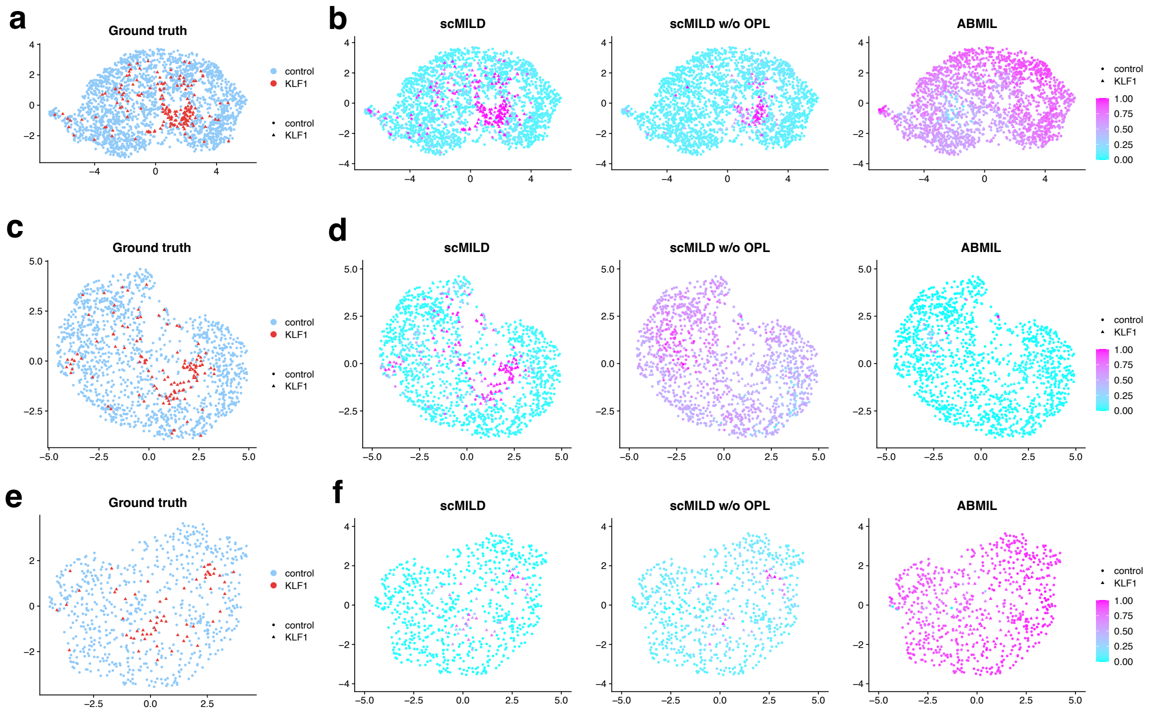


**Supplementary Figure S2.** UMAP visualizations of experiments with lower total cell numbers, illustrating capacity cellular differences under varying conditions (a,b) Experiment with 300 cells per sample; (c,d) with 200 cells per sample; (e,f) with 100 cells per sample, each with 20% perturbed cells. In each set, the left panel shows gorund truth cell labels, with perturbed cells represented as red triangles and control cells as blue circles; the right panel shows cell attention scores from scMILD, scMILD w/o OPL, and ABMIL models.

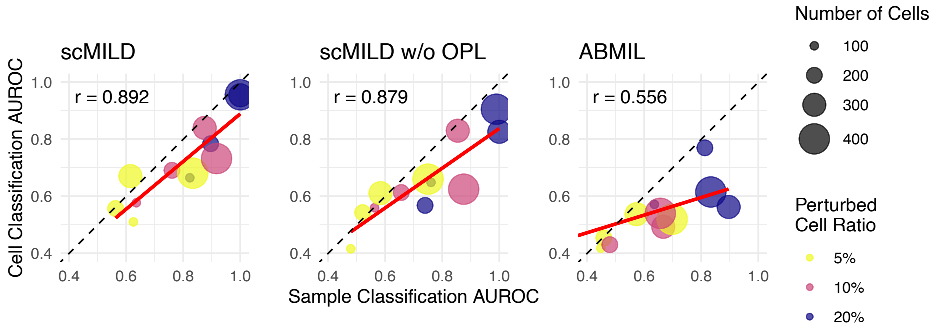


**Supplementary Figure S3.** Scatter plots comparing sample AUROC (x-axis) and cell AUROC (y-axis) for scMILD, scMILD w/o OPL, and ABMIL. Each point represents a simulation setting, with point size indicating the total number of cells per sample and color representing the fraction of perturbed cells.

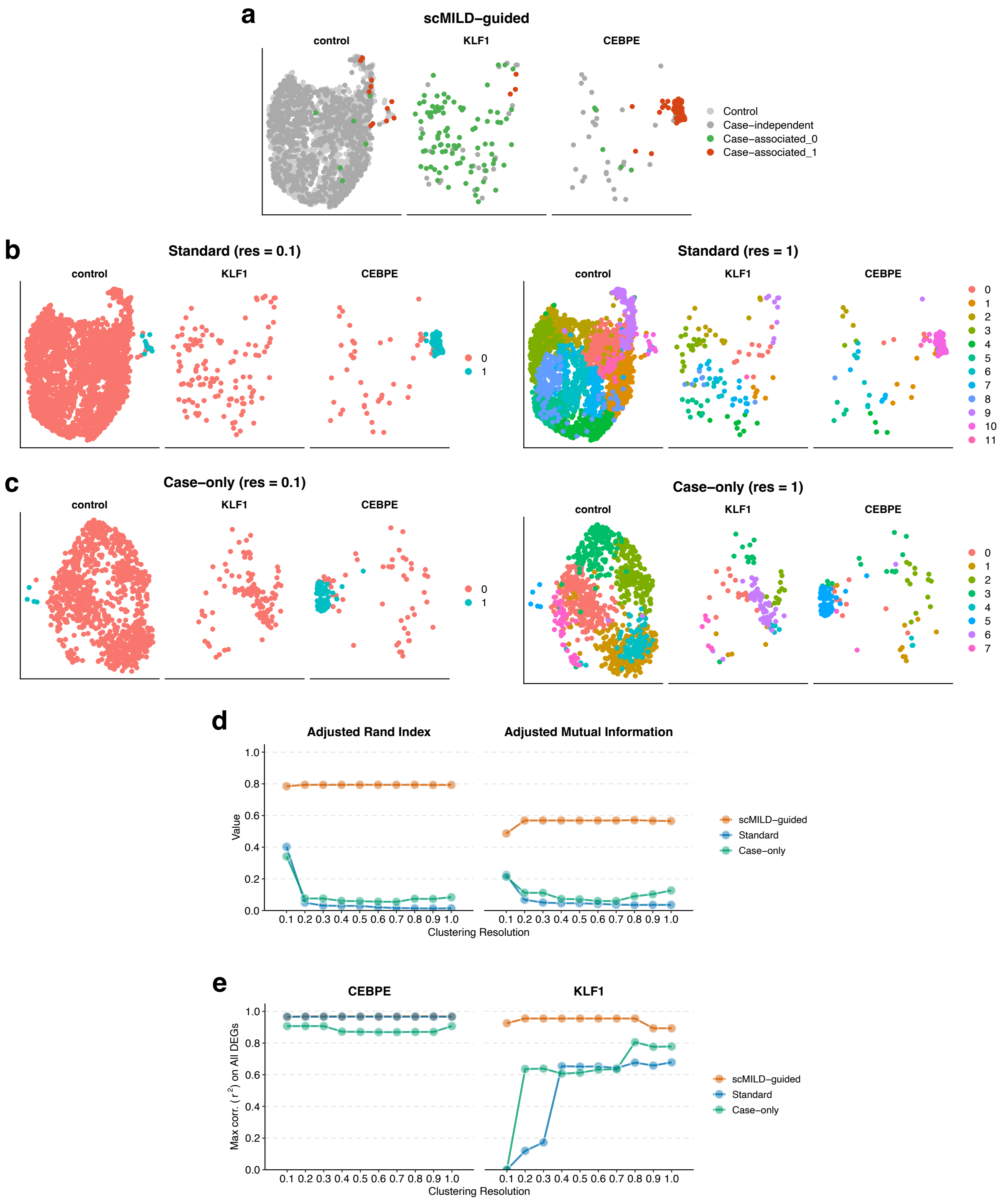


**Supplementary Figure S4.** (a-c) UMAP visualizations of clustering results from (a) scMILD-guided clustering, (b) standard clustering (at resolutions 0.1 and 1.0), and (c) case-only clustering (at resolutions 0.1 and 1.0).
(d) Line plots showing clustering performance metrics (Adjusted Rand Index and Adjusted Mutual Information) across a range of resolution values for the different clustering strategies. (e) Maximum coefficient of determination ($r^2$) from Pearson correlation between the log2 fold-change (log2FC) values from cluster-based DEG analysis and ground truth DEG analysis, calculated using all genes and plotted across a range of resolution values.
